# Frequency coding of multisensory integration in the local field potentials of the medial pulvinar

**DOI:** 10.1101/2024.07.18.604099

**Authors:** Anne-Laure Vittek, Cécile Juan, Corentin Gaillard, Manuel Mercier, Pascal Girard, Suliann Ben Hamed, Céline Cappe

**Author notes:** Corresponding author, Centre de recherche Cerveau et Cognition, CNRS CERCO UMR 5549, Pavillon Baudot, CHU Purpan, 1, place du Dr. Baylac, BP 25202, 31052 Toulouse Cedex.

## Abstract

The pulvinar is a posterior thalamic nucleus, with a heterogeneous anatomo-functional organization. It is divided into four parts, including the medial pulvinar, which is densely connected with primary unisensory and multisensory cortical regions, and subcortical structures, including the superior colliculus. Based on this connectivity, the medial pulvinar may play an important role in sensory processing and multisensory integration. However, its contribution to multisensory integration has rarely been directly investigated. To fill this knowledge gap, two macaque monkeys were trained on a fixation task, during which auditory, visual and audiovisual stimuli were presented. We characterize local field potentials of the medial pulvinar associated with these stimuli. In the temporal domain, we describe an early and a late period showing multisensory integration, both dominated by sub-additive processes (the audiovisual response is inferior to the sum of the unisensory responses). In the frequency domain, multisensory integration, mostly sub-additive, is predominant in the lower frequencies (90% of recorded signals in 4.5-8.5Hz and 96% in 8.5-20Hz). Prevalence largely decreases in high frequencies (54% in 35-60Hz, 37% in 60-120Hz). This suggests that the medial pulvinar is a multisensory hub, integrating visual and auditory information in different frequency bands and contributing to cortico-pulvino-cortical multisensory computational loops.

## Introduction

The pulvinar is a posterior thalamic nucleus, the medial pulvinar being one of its four subdivisions. It is extensively connected with the entire brain. In particular, it is reciprocally connected with unisensory areas (Homman-Ludiye et al. 2020; Froesel et al. 2021) and multisensory areas like the superior temporal sulcus (Jones and Burton 1976; Baleydier and Morel 1992; Romanski et al. 1997; Homman-Ludiye and Bourne 2019), the prefrontal cortex (Romanski et al. 1997; Homman-Ludiye and Bourne 2019; Homman-Ludiye et al. 2020) and the temporal lobe (Trojanowski and Jacobson 1976). Subcortically, the medial pulvinar projects to the amygdala (Jones and Burton 1976; Romanski et al. 1997; Homman-Ludiye and Bourne 2019) and is connected with the superior colliculus (Trojanowski and Jacobson 1975; Benevento and Standage 1983; Homman-Ludiye and Bourne 2019), both of which are multisensory regions. Amongst thalamic nuclei receiving inputs from several modalities, the medial pulvinar is the one with the densest multimodal connectivity and the greatest overlap in synaptic terminals (Cappe, Morel, et al. 2009). These anatomical connections suggest that the pulvinar plays a role in multisensory processing. Numerous single-unit studies have reported sensory responses in the primate pulvinar (Mathers and Rapisardi 1973; Gattass et al. 1978, 1979; Yirmiya and Hocherman 1987; Wilke et al. 2009; Maior et al. 2010; Komura et al. 2013; Nguyen et al. 2013; Van Le et al. 2013, 2014, 2016). Yet, they often included only unimodal stimuli, and were not specifically focused on the medial pulvinar. From a cognitive perspective, the pulvinar is involved in spatial localization (Ward and Arend 2007), visual attention and sensory distractor filtering (Saalmann and Kastner 2009; Snow et al. 2009; Fischer and Whitney 2012), and visually guided movements (Wilke et al. 2010, 2018). Altogether, this has led to the hypothesis that the pulvinar is a modulator of behavioral flexibility (Froesel et al. 2021). Moreover, the pulvinar is thought to participate to coordinated communication between cortical areas (Saalmann and Kastner 2011; Benarroch 2015; Cortes et al. 2020). It is also suggested to be involved in emotion processing (Ward et al. 2005, 2007; Arend et al. 2015). It is proposed that this latter specific function may have appeared under selective evolutionary pressure, such that the pulvinar of primates rapidly detects snakes and other predators (Isbell 2006).

Most neurophysiological recordings in the medial pulvinar have focused on the neuronal properties of this nucleus, thus mostly representing its output (Buzsáki 2004). In the present study, we recorded both single-units and local field potentials (LFP) in the medial pulvinar of two macaques during a fixation task during the presentation of unisensory auditory and visual, and multisensory audiovisual stimuli. The single-unit neuronal data is already reported in Vittek et al. (2023). Specifically, we identified auditory, visual and audiovisual neurons and we demonstrated that multisensory integration in the medial pulvinar is mostly sub-additive and suppressive at the neuronal level. In the following, we focus on medial pulvinar local field potentials, a neurophysiological signal correlating mostly with input signals (Einevoll et al. 2013) and we investigate their multisensory properties in response to audiovisual stimulus presentations. We find massive frequency specific multisensory integration processes across multiple frequency bands in the LFPs of the medial pulvinar, and we characterize their specific multisensory integration properties in both the time and time-frequency domains.

## Materials and methods

### Animals

Two 5-years-old male rhesus monkeys (Macaca mulatta), weighing 7 and 5 kg participated to this experiment. They were naïve to all procedures. They were individually housed during the day (for food and water) and paired at night. Water was given ad libitum, whereas fruits, vegetables and cereals were controlled and given after each training or recording session. The criterion for stopping the experiment was a 10% weight loss. It never happened. All procedures were approved by the National Committee for Ethical Reflection on Animal Testing in compliance with the guidelines of the European Community on Animal Care (authorization number: 01000.02).

### Experimental design and statistical analyses

As in Vittek et al. (2023), monkeys were trained on a fixation task, seated in a primate chair (Crist instruments), head-fixed, in a darkened and sound-attenuated box. They were seated in front of a screen (BenQ, 60 cm diagonal, 1920 × 1080 pixels, 120 Hz), at a distance of 31 cm, with two loudspeakers (Creative Gigaworks t20 serie II) placed at the left and right of the screen. Monkeys were trained five mornings a week. Each correct trial was rewarded with diluted compote (0.05 mL/trial). There were three steps in the task (figure 1.A). First, the monkeys had to fixate a 0.5° fixation point during 500 to 1200 ms. Fixation was controlled thanks to a fixation window defined by a virtual 2×2° square around the fixation point. Second, an auditory, visual or audiovisual stimulus was presented for 250 ms. During this step, the monkey had to maintain central fixation. If fixation was not broken, the monkey was rewarded after a random delay ranging between 300 and 700 ms (third step), otherwise no reward was given. The intertrial interval was 1000 ms. Eye tracking was performed with an eye-tracker (ISCAN ETL 200, Woburn, MA 01801), and the task was controlled with EventIDE software (Okazolab Ltd).

**Figure 1:**
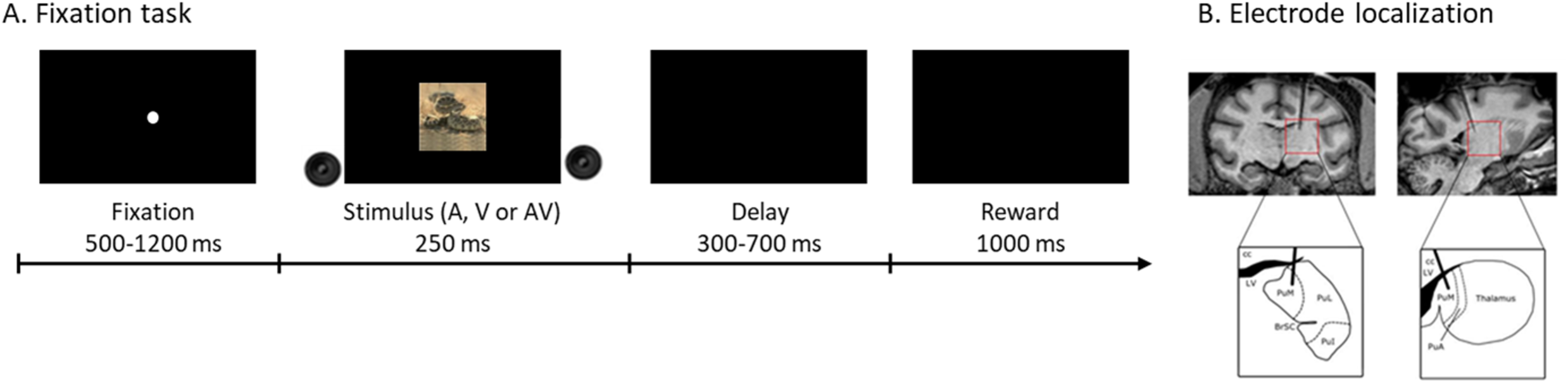
experimental design and recordings localization. **A) Schematic representation of the behavioral task.** The monkey sat in a primate chair in front of a computer screen. The monkey first had to fixate the central point during 500 to 1200 ms. An auditory, visual or audiovisual stimulus was then presented for 250 ms. The monkey had to maintain central fixation during stimulus presentation to get a reward (compote drop). **B) Schematic view showing recordings localizations**. T3 MRI slices of monkey C brain in a frontal (top, left) and a sagittal (top, right) section showing electrode position. Bottom pictures show cerebral regions delimited by the boxes in MRI slices, drawn and labeled according to the Paxinos macaque brain atlas (Paxinos et al. 2000; Saleem and Logothetis 2007). PuM: medial pulvinar, PuL: lateral pulvinar, PuI: inferior pulvinar, PuA: anterior pulvinar, BrSC: superior colliculus brachium, LV: lateral ventricle, cc: corpus callosum.

Three stimuli were used for each modality (nine in total). The auditory stimuli consisted in white noise, a macaque “coo” and a rattlesnake rattle. They were stereo 44-48 kHz waves normalized to 60 dB, with a 3 ms fading-in and fading-out. The visual stimuli were a picture of random dots, a rhesus macaque face picture and a rattlesnake picture. All images were normalized in RGB colors with a color depth of 24, sized 453×453 pixels with a 72×72 dpi resolution (final size 19×19°), with a mean luminance of 120 cd/m², and presented in the center of the screen. The three audiovisual stimuli consisted in the synchronous presentation of an auditory and the congruent visual stimulus (i.e. visual noise with auditory noise, rattlesnake picture with rattlesnake rattle and macaque coo with macaque face). The stimuli were presented around 20 times each, in a random order. The precise onset and offset of visual stimuli presentation was monitored using a photodiode covering a 4×4° white square appearing in the upper right corner of the screen (hidden from the monkey’s view by black tape) at the same time as the stimulus presentation.

### Surgical procedures

Monkeys had two surgeries, one to implant an MRI-compatible headpost (Crist Instrument) to the skull and a second one to implant a footed stainless-steel recording chamber (Crist Instrument) above the right somatosensory cortex (S2) (on average AP = 6.5 and ML = 0.75). Surgeries were described in Vittek et al. (2023). Briefly, surgeries were performed under anesthesia (mixture of tiletamine/zolazepam (Zoletil 50®, 5 mg/kg) and glycopyrrolate bromide (Robinul®, 0.01 mg/kg) for the induction, followed by isoflurane (1.5%) after intubation). Analgesics were administered (Tolfedine 4 mg/kg and buprenorphine chlorhydrate (Vetergesic®) 0.01 mg/kg) during surgery and on the following days, and an antibiotic treatment (Amoxicillin (Clamoxyl® LA, 15 mg/kg)) was administered during the first week. Before the second surgery, the location of the recording chamber was determined by comparing stereotaxic 3T anatomical MRI scans and sections of the stereotaxic atlas of the brain of Macaca mulatta (Paxinos et al. 2000; Saleem and Logothetis 2007). A stereotaxic anatomical MRI was performed after the surgery with a tungsten microelectrode inserted at target coordinates to confirm the locations of the recording sites. The locations of the recording sites are defined relative to the zero coordinates defined in the stereotaxic atlas.

### Recordings

Tungsten microelectrodes (5-7 MΩ at 1 kHz, Frederick Haer Company, Bowdoinham, ME) were inserted daily into the medial pulvinar (figure 1.B) at varying coordinates (ML between 5 and 9 mm, AP between 2 and 7 mm, depth between 17.9 and 23.8 mm below the cortical surface) to record the neuronal activity. Electrode insertion was performed with an oil hydraulic micromanipulator (Narishige MO-972) attached to the recording chamber. The recorded signal was sampled at 40 kHz for neurons and 2 kHz for local field potentials, with a 1401 power acquisition interface (CED, Cambridge, UK) and amplified with a gain of 1000 (NL104) by a Neurolog system (Digitimer, Hertfordshire, UK). A Humbug device (Digitimer) eliminated the 50 Hz noise. The signal was displayed and recorded with Spike2 software (CED).

### Data and statistical analysis

#### Local field potentials in the time domain (evoked potentials)

LFPs recordings were divided into trials of 1.05 s, running from 500 ms prior to stimulus presentation to 550 ms post-stimulus presentation (i.e. covering the 250 stimulus presentation time as well as a 300 ms fixed window into reward expectation time). For each session, trials from the same modality (V: visual; A: auditory; AV: audiovisual) were averaged together. The baseline LFP activity per session and modality was defined as the average LFP activity across the 500 ms fixation window prior to stimulus onset. Kruskal-Wallis tests confirmed, for each LFP recordings, that baselines were not different across conditions (p>0.05). For each trial, LFP in the selected time interval were baseline-corrected by subtracting the baseline from each timepoint, thus re-centering the signals and correcting for potential amplifier slow drift (Mercier et al. 2022).

In order to assess whether the V, A and AV stimuli elicited an evoked response in the LFP signal, Wilcoxon rank sum tests were used to assess whether the post-stimulus evoked response (average LFP activity from 0-550ms following stimulus onset) was significantly different from the response during the baseline epoch (average LFP activity from -500ms to 0ms following stimulus onset). Presence of a significant evoked response was assessed independently for each modality. In order to compare the amplitude of the evoked response in the three V, A and AV modalities, Kruskal-Wallis tests and post-hoc Wilcoxon tests were used. Statistical tests were performed on each timestamp and signals were considered significantly different when p-values were below the 0.05 significance threshold for at least 24 timestamps in 30 successive independent time stamps (i.e. every 15ms as LFP sampling rate = 2 kHz).

As a result of these analyses, LFP signals were classified as non-responsive, auditory, visual or audiovisual. The non-responsive group corresponds to sessions responding to none of the stimuli. The visual group corresponds to sessions responding to visual or to visual and audiovisual stimuli, without any significant difference between the two. Similarly, the auditory group corresponds to sessions responding to auditory or to auditory and audiovisual stimuli, without any significant difference between the two. Finally, the audiovisual group includes all sessions presenting multisensory activity, i.e. i) a response to visual and audiovisual stimuli with a significant difference between the two responses, or ii) a response to auditory and audiovisual stimuli with a significant difference between the two responses, or iii) a response to audiovisual stimuli only, as well as to both visual and auditory stimuli, or iv) a response to all stimuli, in a non-distinctive manner. This analysis was also performed across sessions, considering LFP signal average trace per modality in each session as sample.

This analysis was first performed considering all stimuli irrespective of their identity (noise, macaque and snake) and then independently on each stimulus type. Distributions of sessions’ classification was compared across the three stimuli with a chi-squared test. Statistical thresholds for considering distributions as significantly different was set at p-value<0.05.

#### Local field potentials in the frequency domain

On each independent trial, we applied a continuous wavelet transform on the LFP signals (Oostenveld et al. 2010). Each signal had a total length of 1.05 s, allowing a frequency resolution of 0.5 Hz ranging between 4.5 and 120 Hz, with four cycles for frequencies under 20 Hz and eight cycles for frequencies above 20 Hz. Each trial was then normalized as follows: baseline was subtracted at each timestamp and values were then divided by the baseline, thus correcting for the 1/f power law (Mercier et al. 2022). Four functional frequency bands of interest were determined 4.5-8.5 Hz, 8.5-20 Hz, 35-60 Hz and 60-120 Hz, based on the prevalence of frequency peaks identified in each frequency band (supplementary figure 1). These frequency bands displayed distinct multisensory properties both in the temporal and in the frequency domains (see result section). Power spectra were averaged independently for each of these four frequency bands, and then analyzed separately, similarly to what is done in the time domain. Resultant signals were normalized between 0 and 1. Signals were then smoothed using a boxcar function, which was progressively increased for higher frequencies to compensate for increased temporal resolution in wavelet analysis (width of 7 ms for the 4.5-8.5 Hz frequency band, 15 ms for the 8.5-20 Hz frequency band, and 25 ms for the 35-60 Hz and 60-120 Hz frequency bands).

For individual and across session analyses, sessions were classified, for each frequency band, in the four previously defined groups (non-responsive, auditory, visual, and audiovisual). Statistical tests were performed on each timestamp and signals were considered significantly different when p-values were below the 0.05 significance threshold for at least 80% timestamps in n successive time stamps, n varying across frequency ranges as follows (4.5-8.5 Hz: n=20ms; 8.5-20 Hz: n=50ms; 35-60 Hz and 60-120 Hz: n=75ms).

Similarly to what was performed in the time domain, this analysis was first performed considering all stimuli irrespective of their identity (noise, macaque and snake) and then independently on each stimulus type. Distributions of sessions’ classification was compared across the three stimuli with a chi-squared test. Statistical thresholds for considering distributions as significantly different was set at p-value<0.05.

#### Multisensory integration evaluation

Multisensory integration was evaluated in the time and the frequency domains by computing an additivity index ADI, formulas follows:

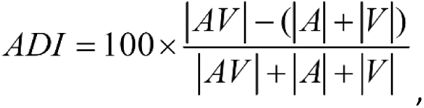

with AV, A, and V the mean audiovisual, auditory and visual responses on specific (identical) time windows, per each multisensory LFP signal. This additivity index ADI allows to define multisensory integration as sub-additive (index<0), additive (index=0) or supra-additive (index>0) (Besle et al. 2004; Kayser et al. 2008).

In the time domain, ADI were computed in two different time windows, corresponding to the two evoked negativities: 30-62 ms (first window), 97-200 ms (second window) (figure 2.A). In the frequency domain, auditory, visual and audiovisual peak power was extracted in each frequency band, on the average of all LFP signals. Then, average power was computed around these peaks (+/-25ms; 4.5-8.5 Hz frequency band: 119-194 ms; 8.5-20 Hz: 28-112 ms; 35-60 Hz: 29-120 ms; 60-120 Hz: 25-129 ms).

**Figure 2:**
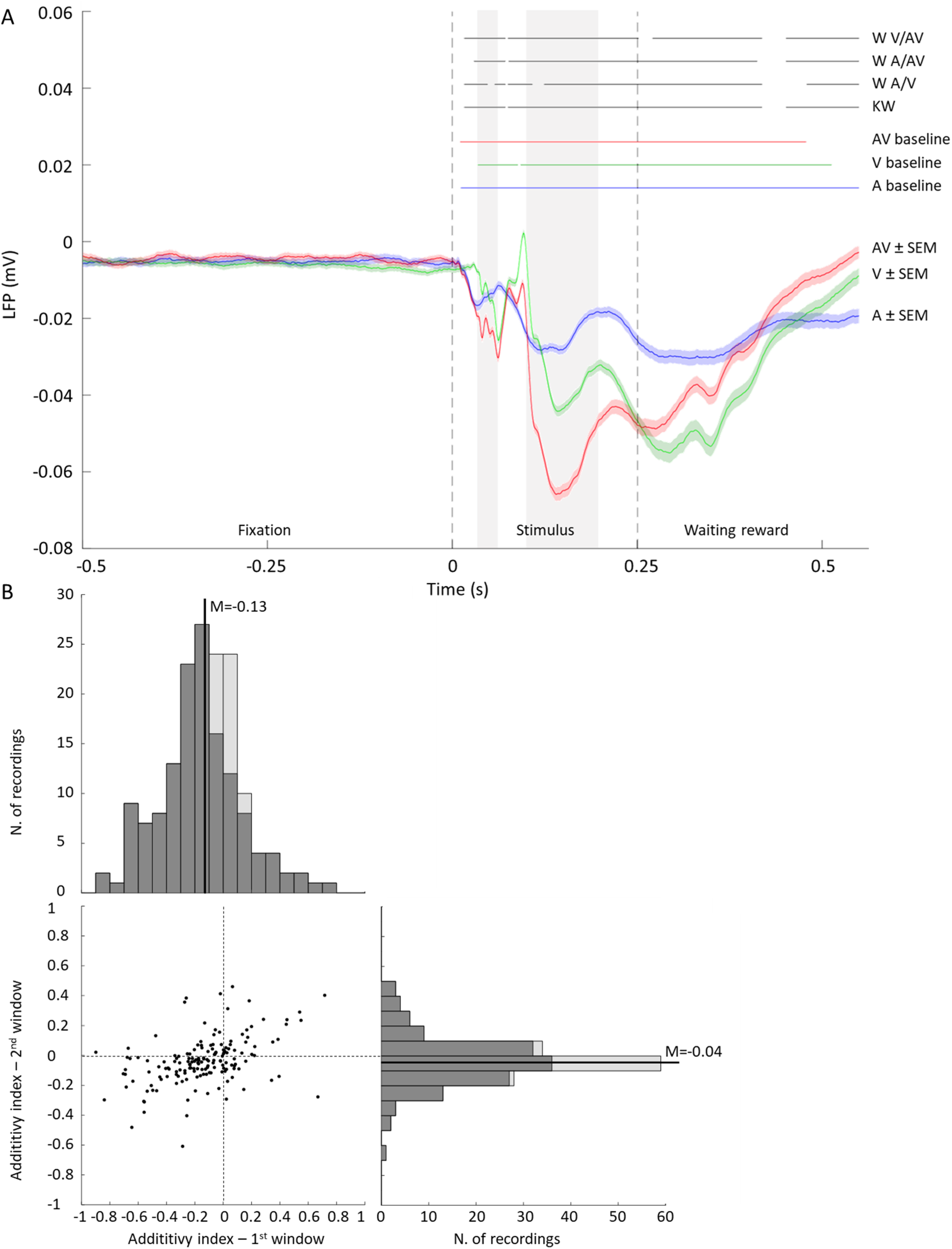
auditory, visual and audiovisual evoked potentials and additivity evaluation. **A) Auditory, visual and audiovisual evoked potentials averaged over all recording sessions (n=163 sessions).** Evoked potentials were calculated over all trials for a given modality (auditory in blue, visual in green and audiovisual in red) with a 3 ms smoothing window. The coloring around evoked potentials corresponds to the Standard Error of the Mean (SEM), using the same color as for evoked potentials. Activity after stimulus onset was compared to baseline (500 ms of fixation before stimulus onset) with a Wilcoxon test, independently for each modality. Colored lines above the graphics indicate timing at which there was a statistical difference (blue: auditory against baseline; green: visual against baseline; red: audiovisual against baseline). Responses to each modality were compared with a Kruskal-Wallis and, when relevant, a paired Wilcoxon test. Timing of statistical differences are indicated with black lines above graphics, in this order: Kruskal-Wallis test assessing statistical differences across modalities (KW, bottom line), Wilcoxon test between auditory and visual responses (W A/V), auditory and audiovisual responses (W A/AV), and visual and audiovisual responses (W V/AV, top line). Gray coloring corresponds to two windows used for computing additivity indexes (30-62 and 97-200 ms). **B) Multisensory integration: additivity index.** Additivity index were computed for each multisensory LFP signal (n=162), on two windows (gray background on figure 2.A), corresponding to the two firsts evoked negativities: 30-62 ms (first window) and 97-200 ms (second window). The additivity index on the first window are plotted as a function of additivity index on the second window (bottom left). Indexes distribution are shown on the top (first window) and on the right (second window). Distribution of additivity index for each window are presented opposite to corresponding axis. Dark gray on the distributions corresponds to index significantly different from 0 (p<0.05), light gray to index not significantly different from 0.

### Data availability

The datasets supporting the current study are available from the corresponding author upon reasonable request.

### Code availability

Analysis were performed on Matlab, with custom codes, available from the corresponding author upon request.

## Results

### Visual, auditory and audiovisual evoked potentials

LFP were recorded in the medial pulvinar (figure 1.B) of two macaques performing a fixation task during the presentation of auditory, visual and audiovisual stimuli (figure 1.A). At the population level (grand-average of all sessions), an evoked potential in response to the auditory, visual and audiovisual conditions can clearly be observed (figure 2.A). Following stimulus onset, these responses are characterized by an initial negativity, followed by a more marked second negativity. These two stimulus-evoked components elicited multisensory properties that were significantly different from the sum of the V and A responses (figure 2.B) (median ± standard error of the median=-0.13 ± 0.03, signtest p<0.001 on the first window, -0.04 ± 0.02, signtest p<0.001 on the second window). A third negativity could yet be observed following stimulus extinction. These three evoked potentials were differently impacted by stimulus sensory modality. The two initial stimulus-ON related evoked potentials were weakest in the auditory modality and highest in the audiovisual condition. In contrast, the stimulus-OFF related evoked potentials were weakest in the auditory modality but highest in the visual condition. This last evoked component elicited sub-additive multisensory properties, in that the AV response was significantly different from each of the unimodal V and A modalities, but additionally significantly smaller than the sum of the unimodal V and A modalities. This is classically taken as a hallmark of multisensory integration. At the individual level, sessions were classified as auditory, visual, audiovisual or non-responsive. Among 163 sessions, 162 were audiovisual (including 156 responding to all stimuli) and only one session was found to be an auditory session. In the time domain, we did not find any uniquely visual or non-responsive session.

### Frequency analysis for auditory, visual and audiovisual responses

#### Prevailing sub-additive multisensory responses

Individual time frequency maps were very diverse. However, functional responses were reliably observed in four different functional frequency bands of interests: 4.5-8.5 Hz, 8.5-20 Hz, 35-60 Hz and 60-120 Hz, as defined based on most frequent structure of individual time frequency maps (figure 3.A, B, supplementary figure S1). The grand-average over all sessions in these four frequency bands are presented in figure 4.A, in response to auditory, visual and audiovisual stimuli. The three responses were significantly different from each other, and on average, AV trials evoked a stronger power in the 4.5-8.5Hz and 8.5-20Hz frequency bands, possibly matching the stronger evoked potential observed for these trials in the temporal domain (figure 2). Please note that in the 4.5-8.5Hz range, the reported frequency represents both the evoked response to the sensory stimuli that is expected to contribute to these low frequencies as well as possibly frequency specific modulations due to theta and alpha long range processes as reported in the cortex (Fiebelkorn et al. 2018, 2019). This was also the case in the higher 60-120Hz frequency band, except that power in the auditory and visual conditions were not significantly different from each other. Although in the 35-60Hz frequency band, AV trials evoked significantly higher power than the V trials, average evoked power on V trials was only slightly weaker than on AV trials. Additivity index ADI revealed sub-additivity at the population level and for all signals. Additivity index were significantly different between the 60-120 Hz frequency band and the other frequency bands again suggesting a possible functional dissociation in multisensory integration between these signals (figure 5). When considering all LFP signals and all frequency bands, all LFP signals but one had at least one multisensory component, and 36.2% of the LFP signals presented multisensory responses in all frequencies. This confirms the strong preponderance of multisensory integration in the medial pulvinar LFP frequency domain, similarly to what has already been described in the time domain.

**Figure 3:**
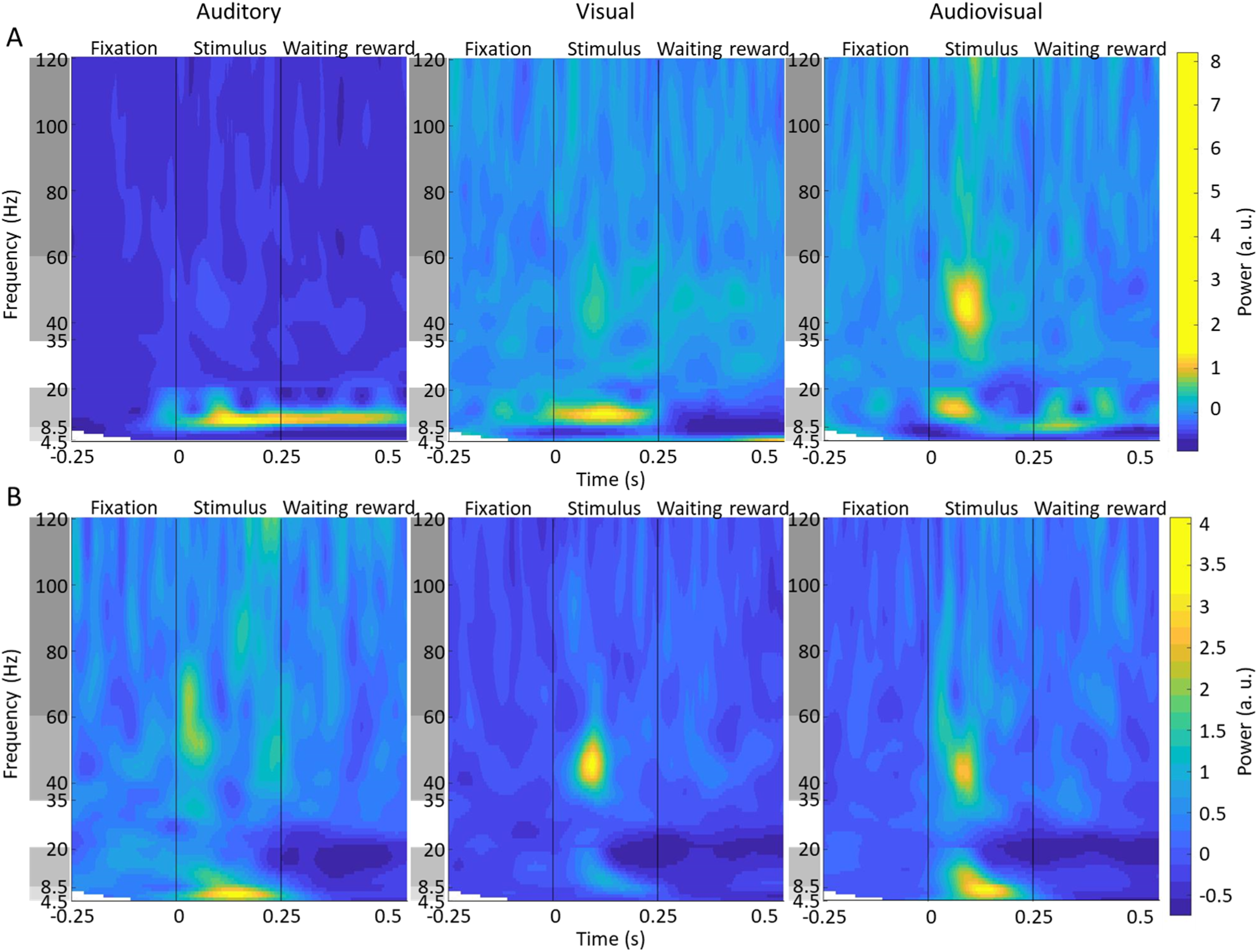
Two example LFP time-frequency responses to visual, auditory and audiovisual stimulations. A) Single LFP signal (LFP#17) time-frequency response to auditory (left), visual (middle) and audiovisual (right) stimuli. B) Single LFP signal (LFP#157) time-frequency response to auditory (left), visual (middle) and audiovisual (right) stimuli. Time-frequency spectra were obtained using wavelet analyses on LFP signals ranging from -0.25 to +0.55 s around stimulus presentation. Frequency ranges of interest are indicated by gray backgrounds (ordinate).

**Figure 4:**
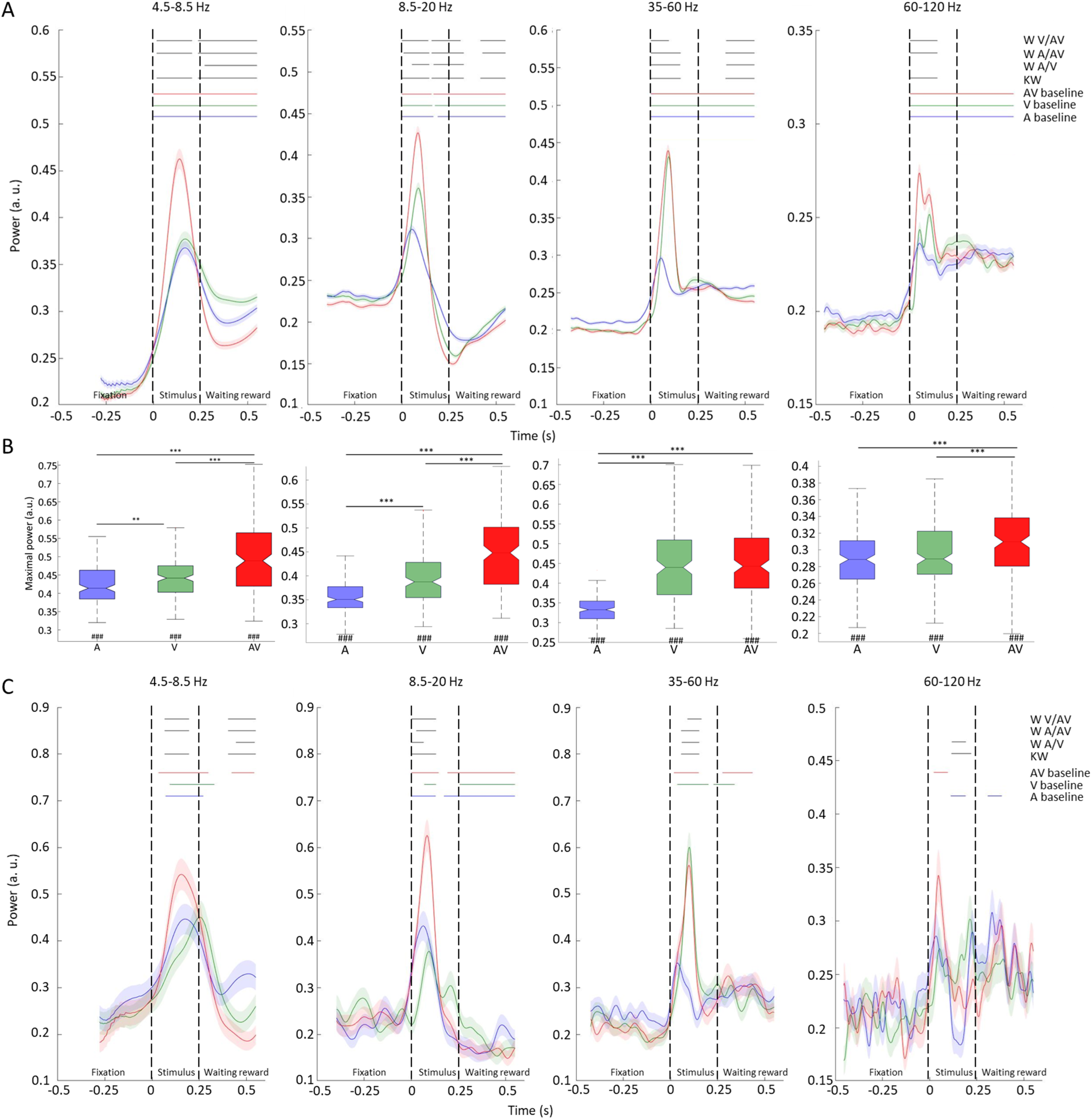
frequency bands average. A) Frequency-band grand-average across all LFP signals, for the auditory (blue), visual (green) and audiovisual (red) stimuli (n=163 sessions). B) Boxplot summarizing distributions of peak power per LFP signal for each frequency band, in response to auditory (A), visual (V), and audiovisual (AV) stimuli. All peak values were different from baseline (Wilcoxon test), sharps below boxplots indicate significance level: ###: P<0.001. Modulation of peak amplitude by the modalities was evaluated by the Kruskal-Wallis test, followed by Wilcoxon paired tests between modalities. Asterisks above boxplots indicate significance level: *: P<0.05, **: P<0.001, and ***: P<0.001. C) Frequency-band averaged activity for the auditory (blue), visual (green) and audiovisual (red) stimuli, for LFP signal #91. Power in time functions were calculated averaging time-frequency power over the four frequency bands of interest (4.5-8.5 Hz, 8.5-20 Hz, 35-60 Hz and 60-120 Hz) following smoothing as described in Material and Methods. For each modality, power after stimulus onset was compared to average power during baseline (500 ms of fixation before stimulus onset) using a Wilcoxon test. Colored lines above the plots indicate timing at which there is statistically significant difference (blue: auditory, green: visual and red: audiovisual). Responses between each modality were compared with a Kruskal-Wallis and, when relevant, paired Wilcoxon test were applied. Timing of statistically significant differences are indicated with black lines above the plots, in the following order: Kruskal-Wallis test (KW, bottom line), Wilcoxon test between auditory and visual responses (W A/V), auditory and audiovisual responses (W A/AV) and visual and audiovisual responses (W V/AV, top line).

**Figure 5:**
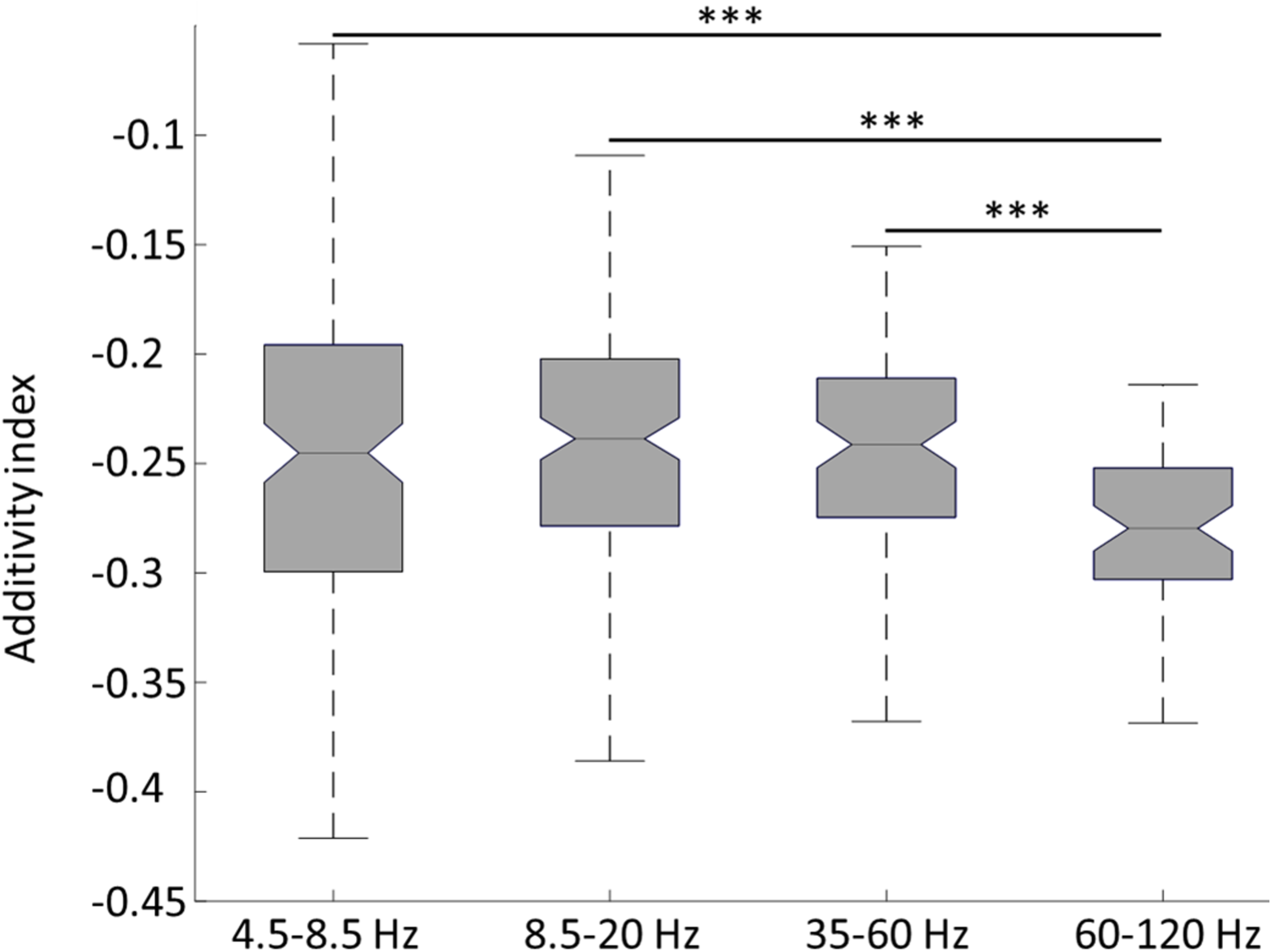
multisensory integration: sub-additivity in all frequencies. Boxplot summarizing distribution of additivity index per LFP signal for each frequency band: data represented as 25^th^ percentile, median and 75^th^ percentile. Multisensory integration was sub-additive in all frequency bands (all signtest, p<0.001), but was different between frequency bands (Kruskal-Wallis test followed by paired Wilcoxon tests). Asterisks above boxplots indicate significance level: ***: P<0.001.

#### Distinct multisensory profiles across frequency bands

Responses were further analyzed at peak values for each frequency band and for all LFP signals. Audiovisual responses at peak were significantly stronger than auditory peak in all frequency bands (figure 4.B). Audiovisual responses at peak were also significantly stronger than visual responses at peak, except in the 35-60 Hz frequency band (figure 4.B). This response pattern can also be seen in individual LFP signals of an exemplar session. The LFP signal presented in figure 4.C responded to all of auditory, visual and audiovisual stimuli in the two lower frequency bands (4.5-8.5 and 8.5-20 Hz). It was thus classified as audiovisual. In addition, evoked power was significantly higher for the AV stimuli than for the visual or auditory stimuli. In the 35-60 Hz frequency band, it responded to visual and audiovisual stimuli, but its auditory response was not significantly different from the baseline. As observed in the grand-average presented in figure 4.A, while the V and AV responses were significantly different, they were very close to each other, suggesting very weak multisensory integration. This LFP signal was again classified as audiovisual in the 35-60 Hz band range. However, in the 60-120 Hz frequency band, evoked power to the auditory and audiovisual stimuli was significantly different from baseline power, but undistinguishable one from the other. This LFP signal was thus classified as auditory for this frequency band. As a result, this LFP signal was considered as audiovisual for the three first frequency bands (4.5-8.5 Hz, 8.5-20 Hz, 35-60 Hz) and unisensory auditory for the last frequency band (60-120 Hz).

Comparing responses between frequency bands revealed a difference between low (4.5-20 Hz) and high (35-120 Hz) frequencies (figure 6 insets). LFP responses in the two lower frequency bands were multisensory (90% of the LFP signals in the 4.5-8.5 Hz frequency band and 96% of the LFP signals in the 8.5-20 Hz frequency band, figure 6). 67% of LFP signals (109) were classified as audiovisual for both frequency bands, and responded to all stimuli. LFP responses in the two higher frequencies were less similar. Indeed, 45 LFP signals were unisensory in one high frequency band and multisensory in the other one. In addition, responses in the higher frequencies were less multisensory. Unimodal responses in the 35-60Hz were dominated by the visual modality, while the higher 60-120Hz was modulated almost equally by auditory and visual cues.

**Figure 6:**
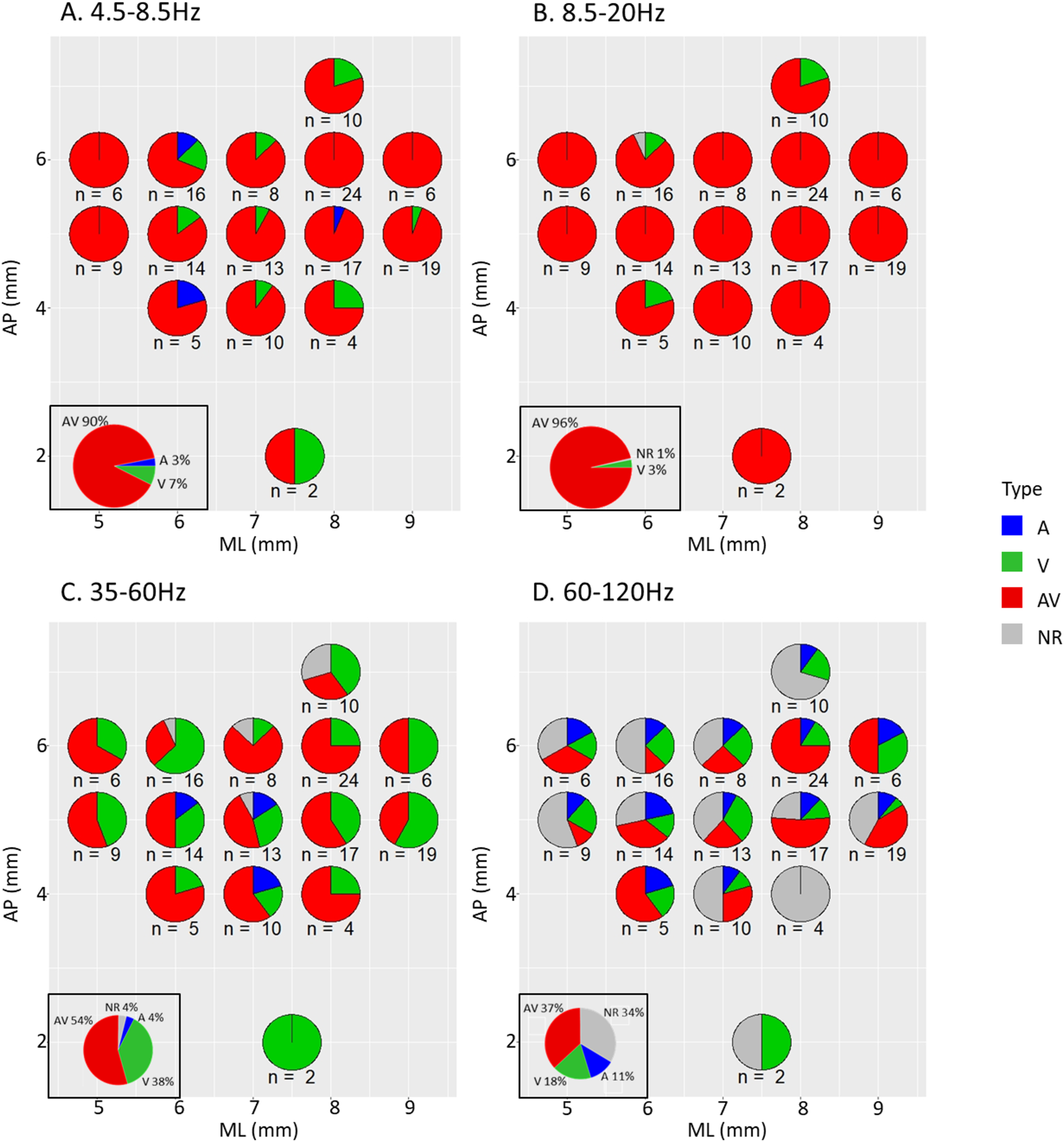
topographic representation of proportions of recording sessions classified by type in each frequency band of interest (n=163 sessions) A-D. Pie charts representing the proportions of sessions classified in respectively 4.5-8.5 Hz (A), 8.5-20 Hz (B), 35-60 Hz (C), 60-120 Hz (D) as auditory only (A, blue) visual only (V, green), audiovisual (AV, red) and non-response (NR, gray), as a function of the anteroposterior (y axis, from 2 to 7 mm) and mediolateral (x axis, from 5 to 9 mm) coordinates of electrode penetration sites. Numbers below pie charts indicates the number of recordings at each site. Boxed pie charts in the low left corner represent proportions on all recording sites together.

We computed for each recoding site and each frequency range the proportion of A, V and AV. Figure 6 represents this information projected onto AP-ML section of medial pulvinar (see also figure 1). While there is a clear difference in the proportion of multisensory signals, decreasing from lower to higher frequency bands, they are prevailing in all frequency ranges. Concerning unisensory signals, visual signals are prevalent in the 35-60 Hz range, while auditory signals are more present in the 60-120 Hz range, leading to a roughly equal proportion of auditory and visual signals in this frequency range. No clear topographical organization can be observed, suggesting that multisensory integration is a core property of medial pulvinar neuronal organization.

### Specificity for stimulus

The observations reported above remained grossly unchanged when analyzing the LFP responses separately for the noise, macaque/coo and rattlesnake/rattle stimuli (table 1). In the time domain, the multisensory LFP signals dominated over unimodal signals (noise: 95.7% of the LFP signals; macaque: 87.7%; snake: 90.2%). In the frequency domain, we again observed different properties between low and high frequencies, with a majority of multisensory sessions in the low frequencies (respectively, for the 4.5-8.5 Hz and the 8.5-20 Hz frequency bands, 79.1% and 87.7% for the noise stimulus, 65.6% and 76.7% for the macaque stimulus, 74.2% and 77.3% for the snake stimulus) and a high proportion of visual sessions in the 35-60 Hz frequency band (29.4% for the noise stimulus, 49.1% for the macaque stimulus, 36.2% for the snake stimulus). Non-responsive LFPs dominated in the highest 60-120 Hz frequency band. The proportion of auditory LFPs was roughly equal to that of the visual, while that of audiovisual LFPs was slightly higher.

**Table 1:** proportion of auditory, visual, audiovisual and non-responsive signals in the time domain and in each frequency bands, for each stimulus, and statistically significant differences between stimuli (chi-squared test) This table indicates the proportion of each type of recordings (in %) for the time domain and for each frequency band in the frequency domain, when each stimulus is examined separately. The statistically significant differences in proportions are indicated in the column “Significant differences” (chi-squared test).

|  |  | Macaque | Noise | Snake | Significant differences |
| --- | --- | --- | --- | --- | --- |
| Time domain | Auditory | 3 | 2 | 1 |  |
|  | Visual | 7 | 2 | 8 |  |
|  | Audiovisual | 88 | 96 | 90 |  |
|  | Non-responsive | 2 | 0 | 1 |  |
| 4.5-8.5 Hz | Auditory | 11 | 9 | 6 |  |
|  | Visual | 15 | 9 | 14 |  |
|  | Audiovisual | 66 | 79 | 74 |  |
|  | Non-responsive | 8 | 3 | 6 |  |
| 8.5-20 Hz | Auditory | 6 | 4 | 2 |  |
|  | Visual | 13 | 6 | 15 |  |
|  | Audiovisual | 77 | 88 | 77 |  |
|  | Non-responsive | 4 | 2 | 6 |  |
| 35-60 Hz | Auditory | 1 | 9 | 7 | Macaque < Noise (P=0.005)<br>Macaque < Snake (P=0.02) |
|  | Visual | 49 | 29 | 36 | Noise < Macaque (P=0.008) |
|  | Audiovisual | 36 | 47 | 26 | Snake < Noise (P=0.005) |
|  | Non-responsive | 14 | 15 | 31 | Snake > Macaque (P=0.003)<br>Snake > Noise (P=0.007) |
| 60-120 Hz | Auditory | 4 | 10 | 7 |  |
|  | Visual | 11 | 13 | 10 |  |
|  | Audiovisual | 17 | 23 | 11 | Noise > Snake (P=0.01) |
|  | Non-responsive | 68 | 54 | 72 |  |

Statistically, there were some differences between the three stimuli. These differences were specific to the two higher frequency domains and distinct for each frequency domain. In the 35-60 Hz frequency band, there were less auditory LFP signals responsive to the macaque vocalizations (1% of the signals) than to the noise (9%) stimulus (chi-squared test, p=0.005) and to snake (7%) stimulus (p=0.02). There were less visual signals responsive to noise (29%) compared to the macaque (49%) stimulus (p=0.008). There were less multisensory signals responsive to the snake (26%) stimulus than to the noise (47%) stimulus (p=0.005). There were also more unresponsive signals to the snake (31%) stimulus than to the macaque (14%) stimulus (p=0.003) and to the noise (15%) stimulus (p=0.007). In the 60-120 Hz frequency band, we observed more multisensory signals responsive to the noise (23%) than to the snake (11%) stimulus (p=0.01). This hints for a differential coding of visual and auditory stimuli in the different frequency bands.

## Discussion

LFPs were recorded in the medial pulvinar of macaques performing a fixation task during the presentation of auditory, visual and audiovisual stimuli. Sensory responses were characterized in the time and frequency domains at the population level. Multisensory integration was identified in an early and a later window during stimulus presentation. Responses of individual LFPs depended on the frequency band, suggesting a frequency coding for multisensory integration in the medial pulvinar.

### Unisensory and multisensory inputs to the medial pulvinar

Most studies about sensory processing in the pulvinar have relied on single units (Mathers and Rapisardi 1973; Gattass et al. 1979; Benevento and Miller 1981; Yirmiya and Hocherman 1987; Komura et al. 2013), and mainly unisensory stimulations (but see Gattass et al., 1978; Avanzini et al., 1980; Vittek et al., 2023). Only few studies have analyzed LFP properties in the pulvinar, and were not specific to sensory perception: they covered attention (Zhou et al. 2016; Fiebelkorn et al. 2019) and perceptual suppression (Wilke et al. 2009), all focusing on the visual modality. LFPs recorded in any cortical or subcortical region globally correspond to inputs to this region (Pesaran 2010; Einevoll et al. 2013) whereas single-units correspond to the output (Buzsáki 2004). Thus, our study is the first to show that the medial pulvinar receives auditory and audiovisual inputs bringing functional support to the auditory and multisensory fMRI activations described in macaque medial pulvinar (Froesel et al. 2022a; Froesel et al 2024a) as well as to the dense functional connectivity of the pulvinar with multisensory associative cortices (Froesel et al., 2024b). It also confirms that the medial pulvinar receives dense visual inputs. We further show a high prevalence of multisensory LFPs. This high proportion of multisensory inputs contrasts with that of the multisensory outputs (spikes): 99% of our LFPs are identified as multisensory, whereas only 42% of the single-units recorded simultaneously were multisensory (Vittek et al. 2023). This suggests that the internal computations within the medial pulvinar segregate pervasive multisensory input into independent sensory channels, possibly supporting behavioral and cognitive flexibility.

### Frequency coding of sensory information

Time-frequency analyses revealed that single LFPs were frequency specific. In particular, while multisensory integration was sub-additive in all frequencies, it was significantly more sub-additive in the 60-120 Hz frequency band. We propose that sensory (unisensory and multisensory) information is multiplexed in different functional frequency bands, each band potentially carrying different types of information, possibly due to different groups of cells contributing to each process. These signals could project onto the entire medial pulvinar population, or alternatively, onto distinct pulvinar subpopulations, possibly supporting specific pulvino-cortical loops (Froesel et al., 2024). Such dissociated frequency coding has been proposed in the context of unisensory information. LFPs were recorded in V1 of anesthetized macaques, during unisensory visual movie presentations (Belitski et al. 2008, 2010; Mazzoni et al. 2013) and in the auditory cortex of awake macaques passively listening to sounds (Belitski et al. 2010). Mutual information was computed across LFPs, revealing two informative frequency bands, in low and high frequencies without significant redundancy. Likewise, recording LFPs in the gustatory cortex of awake rats, during tastants and water delivery on the tongue, Pavão and collaborators (Pavão et al. 2014) proposed that different frequency channels are used to code multiple features of a stimulus. More recently, LFPs were recorded in epileptic humans (Sabra et al. 2020) while presenting images of three categories. With principal component and decoding analyses, the authors concluded that different kinds of visual information can be encoded in the power of different spectral components.

Thus, unisensory studies in different species present data showing different information encoded in different LFP frequency bands. We would like to extend these results to multisensory information and to the medial pulvinar. Specifically, we propose that different sensory information are carried by different frequency bands. In this hypothesis, one signal (the LFP) would provide multiple information about one or more sensory modalities, by separating the information on different channels (the frequencies). The functional specificity of each encoding frequency channel still has to be investigated.

### Frequency dependent multisensory integration processes

In the cortex, low frequencies (around 8-15 Hz) are associated with feedback signals, while higher frequencies (around 30-80 Hz) are associated with feedforward signals and local spiking activity (Bastos et al. 2014; van Kerkoerle et al. 2014; Jensen et al. 2015; Michalareas et al. 2016). It is unclear how this generalizes from the cortex to subcortical structures. Medial pulvinar LFPs were modulated, in the high frequencies, by all types of sensory stimuli. This suggests that the medial pulvinar receives unisensory and multisensory feedforward signals. This is consistent with the anatomical connections of the medial pulvinar, receiving afferences from unisensory and multisensory cerebral structures. Auditory responses were less frequent relative to visual and audiovisual responses in these high frequency ranges. This could be due to the fact that gamma activity magnitude varies with task relevance (van Kerkoerle et al. 2014). None of auditory and visual information was relevant. However, in diurnal primates, visual information is automatically granted relevance, thus accounting for the prevalence of visual responses.

In the lower frequencies, most LFPs responded to all sensory stimuli. Audiovisual responses were sub-additive indicating an active multisensory integration process. This suggests that the medial pulvinar receives from the cortex an integrated multisensory input. Functional coupling between the medial pulvinar and the cortex (prefrontal area FEF and parietal area LIP), has been described in these frequency bands during attention orientation (Fiebelkorn et al. 2019). We predict that similar functional coupling will also be at play between the medial pulvinar and specific cortical regions during multisensory integration. For example, the active multisensory integration patterns of audiovisual social stimuli recently described (Froesel et al. 2022a) are very similar to those described in the same task in the superior temporal sulcus (Froesel et al. 2022b), suggesting a functional link between these two regions. It has been proposed that the complex recurrent cortico-pulvino-cortical connectivity might play a crucial role in organizing cortical intra- and inter-areal activity (Lakatos et al. 2016, Froesel et al., 2024). This has been explored in the context of attentional regulation (Kastner et al., 2020 for a review). Similar approaches will have to be applied to multisensory integration to clarify the role of cortico-pulvino-cortical connectivity in this context.

### The multiple sensory functions of the medial pulvinar

While the specific role of the pulvinar is still under investigation, cumulative evidence allows to propose a set of functions (Froesel et al. 2021 for review). A first known role of the pulvinar is a sensory role. Anatomically, the medial pulvinar has dense sensory connections (Cappe et al. 2007; Cappe, Rouiller, et al. 2009). Among all thalamic nuclei presenting overlapping territories of projections from different sensory cortices, the medial pulvinar is the largest one (Cappe, Rouiller, et al. 2009). Electrophysiological studies have shown sensory responses in the pulvinar and specifically in the medial pulvinar (Mathers and Rapisardi 1973; Yirmiya and Hocherman 1987; Maior et al. 2010; Van Le et al. 2013, 2014, 2016; Vittek et al., 2023). The present work refines our understanding of the sensory role of the medial pulvinar, showing that, in addition to sending sensory information to connected brain regions through spiking activity, it receives unisensory and multisensory information, as measured through LFPs, and further processes this information, thus contributing to multisensory integration. A second proposed role for the pulvinar, linked to the first one, is its involvement in sensory distractor filtering (Fischer and Whitney 2012) and visual selection and attention (Saalmann and Kastner 2009). This role, relying on sensory perception, allows to detect important stimuli and respond rapidly and adaptively to them. In our study, all stimuli and modalities were equally important so it is difficult to infer any specific mechanism about distractor filtering. However, in a similar experiment in which the monkey would have to respond to only one modality, we expect a larger proportion of unisensory signals in response to this modality, at least in high frequencies, as these frequencies are modulated by task relevance (van Kerkoerle et al. 2014). Another role of the medial pulvinar concerns emotion processing (Ward et al., 2007; Arend et al., 2015 for review). Accordingly, the medial pulvinar is proposed to be part of two emotion processing pathways, with the amygdala and the superior colliculus. With these three roles taken together (sensory perception, visual selection and attention, emotion processing), the pulvinar is expected to contribute to the selection of sensory inputs and modulate behavioral flexibility (Froesel et al. 2021).

Finally, the pulvinar is probably involved in the communication between cortical areas, by modulating the signal transfer between them (Saalmann and Kastner 2011; Benarroch 2015) and improving the signal-to-noise ratio of transmitted information, by modulating the synchrony between these areas (Fries 2015). Indeed, according to the replication principle, all directly linked cortical areas would also be indirectly connected via the pulvinar (Sherman 2016, 2017). These cortico-pulvino-cortical pathways would be faster than cortico-cortical projections (Sherman and Guillery 2002; Cappe, Morel, et al. 2009).

To conclude, our study demonstrates that the pulvinar receives unisensory and multisensory information. This information, multiplexed in different frequency bands, is central to the already proposed pulvinar functions. Future studies using multi-site recordings of single-units and LFP simultaneously, in the pulvinar and in the cortex would allow to investigate how these findings articulate with our current understanding of multisensory integration in the cortex. This would contribute to a better understanding of the functions of the pulvinar.

## Acknowledgments

This work was supported by the French National Research Agency (ANR) ANR IBM 12PDOC-0008-01 and ANR-16-CE37-0009 NeuroCim. M.M. is now affiliated to INSERM, INS UMR 1106, Institut de Neurosciences des Systèmes, Aix Marseille Université, Marseille, 13005 France.

## Declaration of interests

The authors declare no competing interests.

## Supplementary material

**Supplementary Figure S1:**
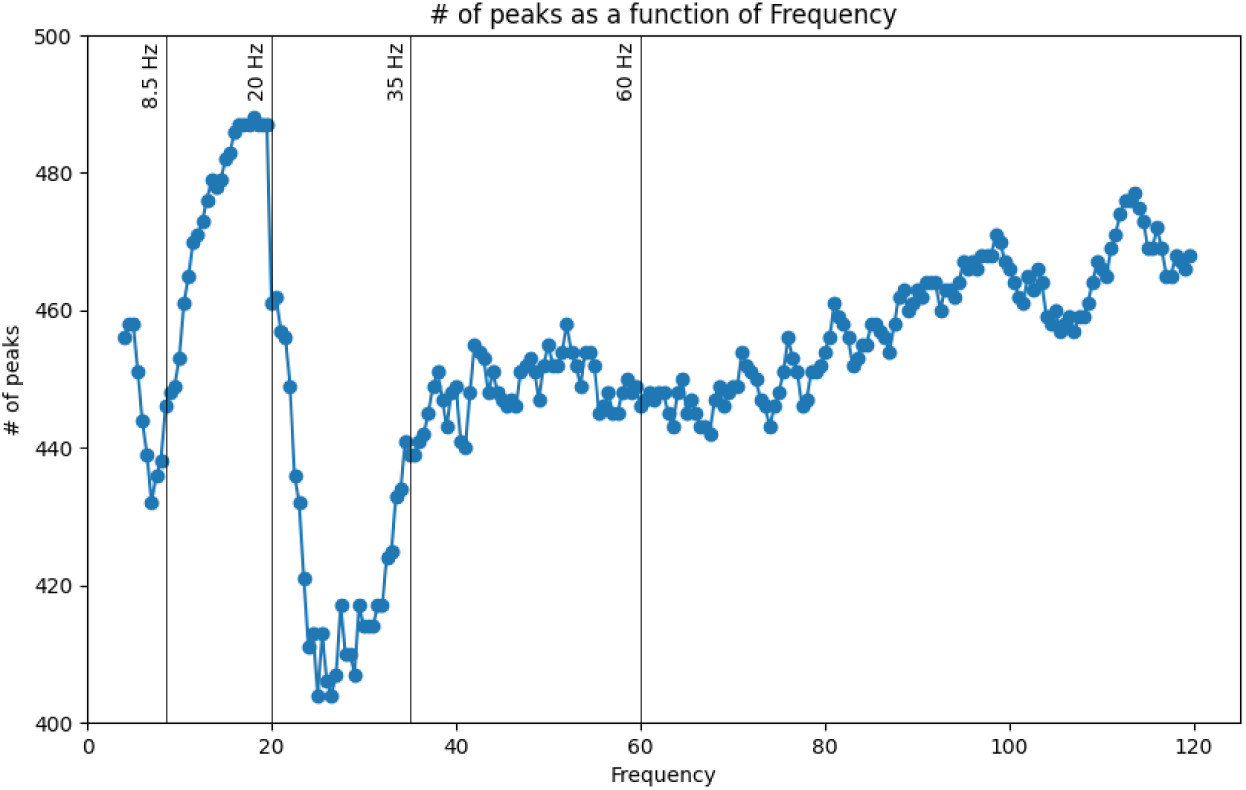
Distribution of number of peaks as a function of frequency and characterization of the different frequency bands on which the analyses were performed (4.5-8.5 Hz, 8.5-20Hz, 35-60 Hz, 60-120 Hz). These frequency bands displayed distinct multisensory properties both in the temporal and in the frequency domains (see result section).

## Notes

### Competing Interest Statement

The authors have declared no competing interest.

